# Deep Learning and Transfer Learning for Brain Tumor Detection and Classification

**DOI:** 10.1101/2023.04.10.536226

**Authors:** Faris Rustom, Pedram Parva, Haluk Ogmen, Arash Yazdanbakhsh

**Affiliations:** Computational Neuroscience and Vision Lab, Neuroscience Program, Boston University; Department of Radiology, VA Boston Healthcare System; Boston University Chobanian & Avedisian School of Medicine; Harvard Medical School; Department of Electrical & Computer Engineering, Laboratory of Perceptual & Cognitive Dynamics, University of Denver; Department of Psychological and Brain Sciences, Computational Neuroscience and Vision Lab, Center for Systems Neuroscience, and Program for Neuroscience, Boston University

## Abstract

Convolutional neural networks (CNNs) are powerful tools that can be trained on image classification tasks and share many structural and functional similarities with biological visual systems and mechanisms of learning. In addition to serving as a model of biological systems, CNNs possess the convenient feature of transfer learning where a network trained on one task may be repurposed for training on another, potentially unrelated, task. In this retrospective study of public domain MRI data, we investigate the ability of neural network models to be trained on brain cancer imaging data while introducing a unique camouflage animal detection transfer learning step as a means of enhancing the network’s tumor detection ability. Training on glioma and normal brain MRI data, post-contrast T1-weighted and T2-weighted, we demonstrate the potential success of this training strategy for improving neural network classification accuracy. Qualitative metrics such as feature space and DeepDreamImage analysis of the internal states of trained models were also employed, which show improved generalization ability by the models following camouflage animal transfer learning. Image sensitivity functions further this investigation by allowing us to visualize the most salient image regions from a network’s perspective while learning. Such methods demonstrate that the networks not only ‘look’ at the tumor itself when deciding, but also at the impact on the surrounding tissue in terms of compressions and midline shifts. These results suggest an approach to brain tumor MRIs that is comparatively similar to that of trained radiologists while also exhibiting a high sensitivity to subtle structural changes resulting from the presence of a tumor. These findings present an opportunity for further research and potential use in a clinical setting.

## Introduction

### Convolutional Neural Networks and Transfer Learning

Sustained and recent progress in Artificial Intelligence (AI) within last decades has been reaching out to many subfields in medicine including radiology and imaging-based inferences [1]–[4]. Convolutional neural networks (CNNs) are powerful tools that are iteratively trained on large image datasets for object recognition and classification tasks. CNNs learn to extract salient features pertaining to object classes and associate these features with specific category labels [5]. In doing so, a trained CNN learns to detect these same features in previously unseen testing images, and thus classifies them under the appropriate training label accordingly. CNNs also have the capacity for “transfer learning,” a training paradigm where a model trained on one task can be repurposed for a new, related task using a different dataset while still using the trained weights from the original task [6]–[8]. This method typically confers an advantage when the original weights prove beneficial while training on the task at hand. Image-based tumor classification involves a difficult pattern recognition problem and can therefore benefit from a transfer learning approach.

### Transfer Learning from Camouflage Animal Detection Task

Although camouflage animal detection and brain tumor classification tasks involve different images, there might be a parallel between an animal hiding through natural camouflage and a bundle of cancerous cells blending in with the surrounding healthy tissue. The learned process of generalization - the grouping of different appearances under the same object identity - is essential to a network’s ability to detect camouflaged objects and is why such training could prove advantageous for a tumor detection task [9]. A trained CNN’s ability to generalize would be inherited through transfer learning from camouflage animal detection to improve its performance on tumor detection and classification [10].

### Feature Spaces as a Metric for Neural Networks’ Internal Representation

Testing accuracy does not describe the changes in internal representation of data by the network in response to training, nor does it directly demonstrate a network’s generalization ability. The activity of thousands of neurons (referred to as nodes) within a CNN can be extracted after training to generate a feature space showing the internal representation of stimuli by the network [11]. Feature spaces present the network’s distribution of every test image relative to one another, providing valuable insight into the network’s organization of data following training. The relationship between individual data points in such a space can be analyzed through the Universal Law of Generalization, which states that the perceived likeliness of two images increases as the distance between them in a feature space decreases [12].

## Methods

### Building Datasets for Neural Network Training

CNNs were trained on a brain tumor classification task using post-contrast T1-weighted and T2-weighted astrocytoma, oligoastrocytoma, oligodendroglioma, and normal (non-cancerous) MRIs obtained from two sources. The majority of our glioma MRIs were obtained from public online repositories such as Kaggle and the Cancer Imaging Archive (TCIA) [13]. For normal MRIs, we used deidentified brain MRIs that was provided to us by co-author Dr. Pedram Parva from the Boston VA Healthcare System approved by Department Of Veterans Affairs, VA Boston Healthcare System R&D Committee (Protocol number: [1628677-1]) . This source accounted for all normal MRIs, and supplemented our existing glioma dataset. All sources used gathered data under HIPAA compliance, and a waiver of HIPAA authorization was approved by the IRB on 4/04/2022 for the Boston VA Healthcare System data since all provided data was anonymized and deidentified. Age and sex of subjects was not available in public datasets used here, and are thus not included in the study. Manual pre-processing of these images resulted in 264 viable axial glioma MRIs (73 astrocytomas, 44 oligodendrogliomas, and 27 mixed tumors formerly described as oligoastrocytomas, and 120 normal).

The camouflage animal datasets used here were imported from a previous project in which neural networks were trained for the detection of camouflage animals [10]. It consisted of almost 3,000 images of clear and camouflaged animals divided into 15 categories.

### Neural Network Training Parameters

AlexNet is a convolutional neural network consisting of a 25-layer series architecture, imported via MATLAB version 2024a. AlexNet is pretrained on over a million images in up to 1000 object categories, and was used as the basis of the trained networks described below*. (*Technically, all models described here are transfer trained since we are using pretrained AlexNet weights at baseline. However, in this paper we are using the term transfer learning to describe models that have had an additional sequence of transfer learning beyond its initial baseline). All images were resized to 227×227 pixels to fit the AlexNet input requirements and were randomly split into training (70%) and testing (30%) sets patient-wise, such that images from the same patient were not cross-contaminated between sets to avoid overfitting. Training was performed on the same machine with the same parameters for all models. Modeling and data analysis code can be accessed with the following link: https://github.com/frustom/Glioma-classification (ID: 4809128)

### Training Neural Networks on Datasets

Two neural networks were trained in this project on the detection and classification of post-contrast T1 and T2 MRIs: T1Net and T2Net, respectively. The dataset included astrocytomas, oligoastrocytomas, and oligodendrogliomas as the tumor categories, and normal brain MRIs as a control. Although we recognize the outdated oligoastrocytoma label [14], we included the images since they were readily available in the datasets used and thus serve as an additional challenge for the neural network to classify appropriately with respect to the other glioma classes.

The previously mentioned camouflage animal detection networks were used as seeds for the tumor classification networks ExpT1Net and ExpT2Net, which both used the ExpCamoNet layer weights as a starting point for training.

### Dimensionality Reduction

To map the feature spaces (examples in Figs 1&2), the vector of activation values of each category node was extracted from the last Fully Connected (FC) layer of a CNN. This vector contains the activation values of every testing image across all category nodes. Principal Component Analysis (PCA) was used as a dimensionality-reduction method to simplify the larger activation matrices to three key dimensions that are representative of the overall activation values. The first three Principal Components were then plotted to model the networks’ feature space in three dimensions. In these feature spaces, each point represents the activation of a single testing image on the trained FC layer nodes and its position within the feature space, as part of a color-coded region, is a visualization of the networks’ internal representation of the data.

**Figure 1.**
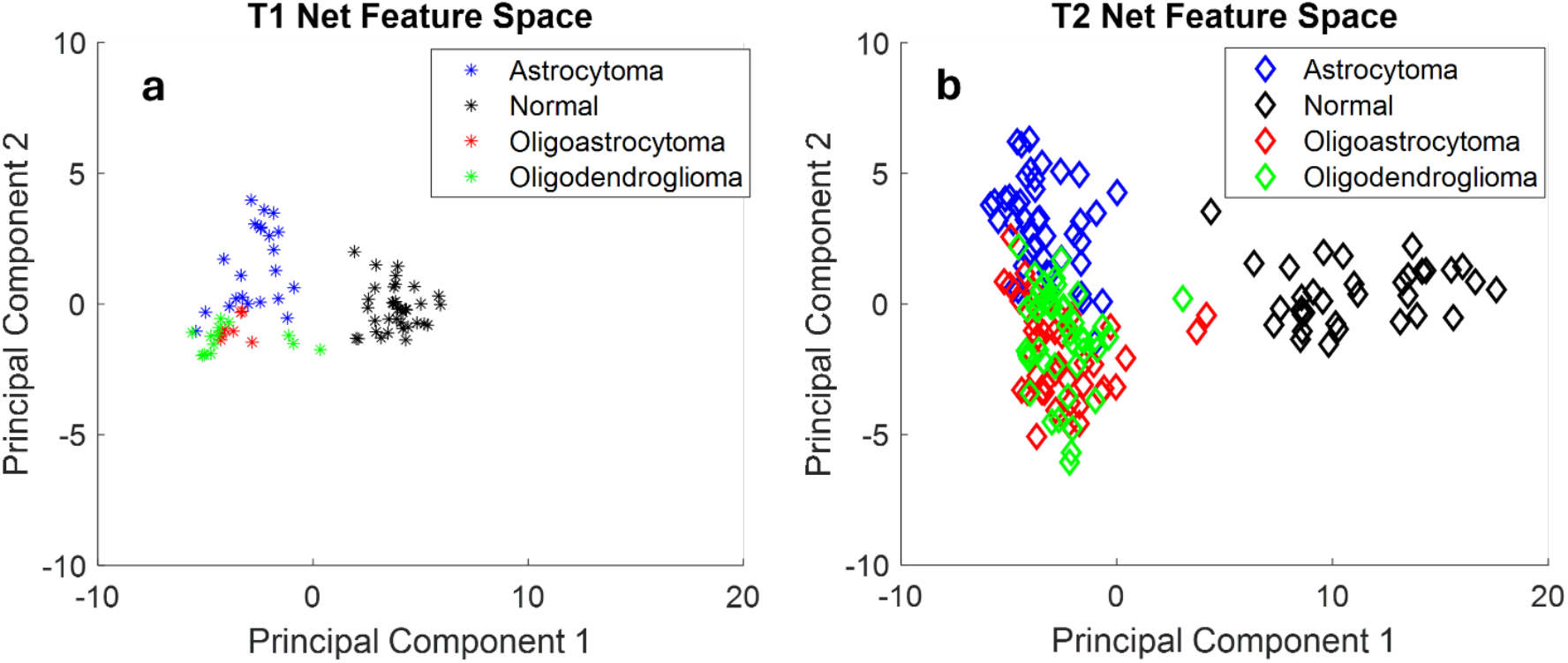
Feature spaces for T1Net (**a**) and T2Net (**b**) across first two Principal Components.

**Figure 2.**
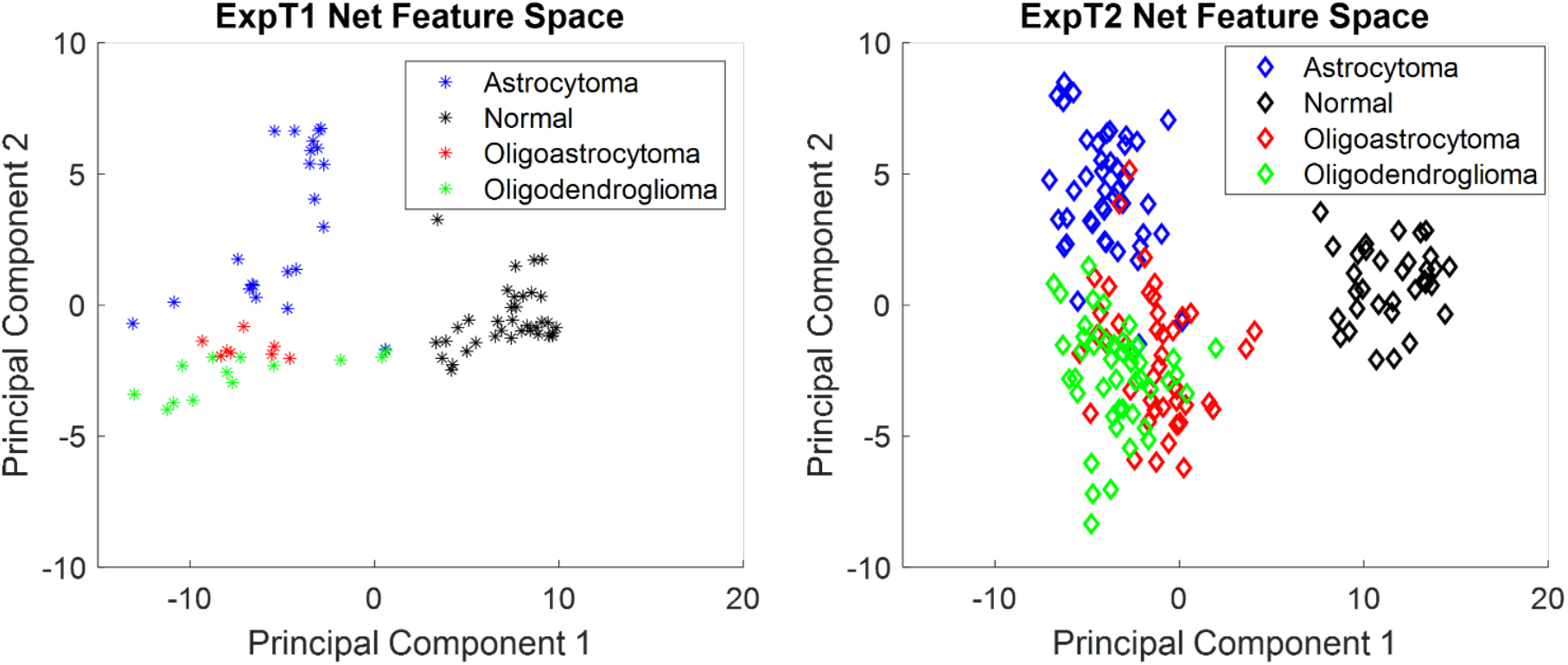
Feature spaces for ExpTINet (**a**) and ExpT2Net (**b**) across first two Principal Components.

### DeepDreamImage

DeepDreamImage (DDI) is a feature visualization technique that offers an alternative method of visualizing the internal state of a trained CNN [15]. This gradient ascent (as opposed to gradient descent) algorithm passes a given image through the network, and calculates the derivative of the image with respect to the activation values of the chosen layer. The algorithm seeks to maximize the loss, which is the sum of the activations in that layer, for the image to increase the activation of that layer. The gradients of this new ‘excited’ image are calculated with respect to the original image, and then added to the original image. This enhances the patterns recognized by the network, and the result is an image that maximally activates the specific category node at the chosen layer. In other words, this algorithm generates a visual ‘prototype’ for each object category resembling the model’s internal representation of that category.

### Image Sensitivity Maps

To determine which features of a given image were most important for the neural networks decision making after training, image sensitivity maps were generated to visualize feature saliency. Both the gradCAM (gC) and occlusionSensitivity (oS) MATLAB functions were used to generate heat maps overlaying select representative images of each training category with an associated color bar representing the intensity of the network’s sensitivity to a given area. The oS function iteratively occludes small areas of the image and calculates the change in probability score for a given class as the occluding object moves across the entire image [16], whereas the gC function uses classification score gradients with respect to convolutional features to determine which features are most crucial for classification [17]. We applied these functions on activation values in both the softmax and fully connected layers to generate image sensitivity maps for both cancerous and normal subjects on all trained neural networks.

## Results

### T1Net, T2Net Performance and Feature Space

Both T1Net and T2Net showed a modest performance on the glioma classification task. T1Net had an average accuracy of 85.99%, with T2Net trailing behind at 83.85%. Table 1 outlines the accuracies and statistics of each trained network. T1Net and T2Net both had near perfect accuracies on normal brain images, with only 1-2 false negatives between both networks, demonstrating a strong ability to differentiate between cancerous and normal brains. The networks struggled more with glioma subtype classification, with each network showing contrasting performance on glioma categories. T1Net’s best glioma category was astrocytoma (95.46%) and its worst was oligoastrocytoma (12.50%), whereas T2Net’s best category was oligoastrocytoma (93.33%) and its worst was astrocytoma (74.42%).

**Table 1.**
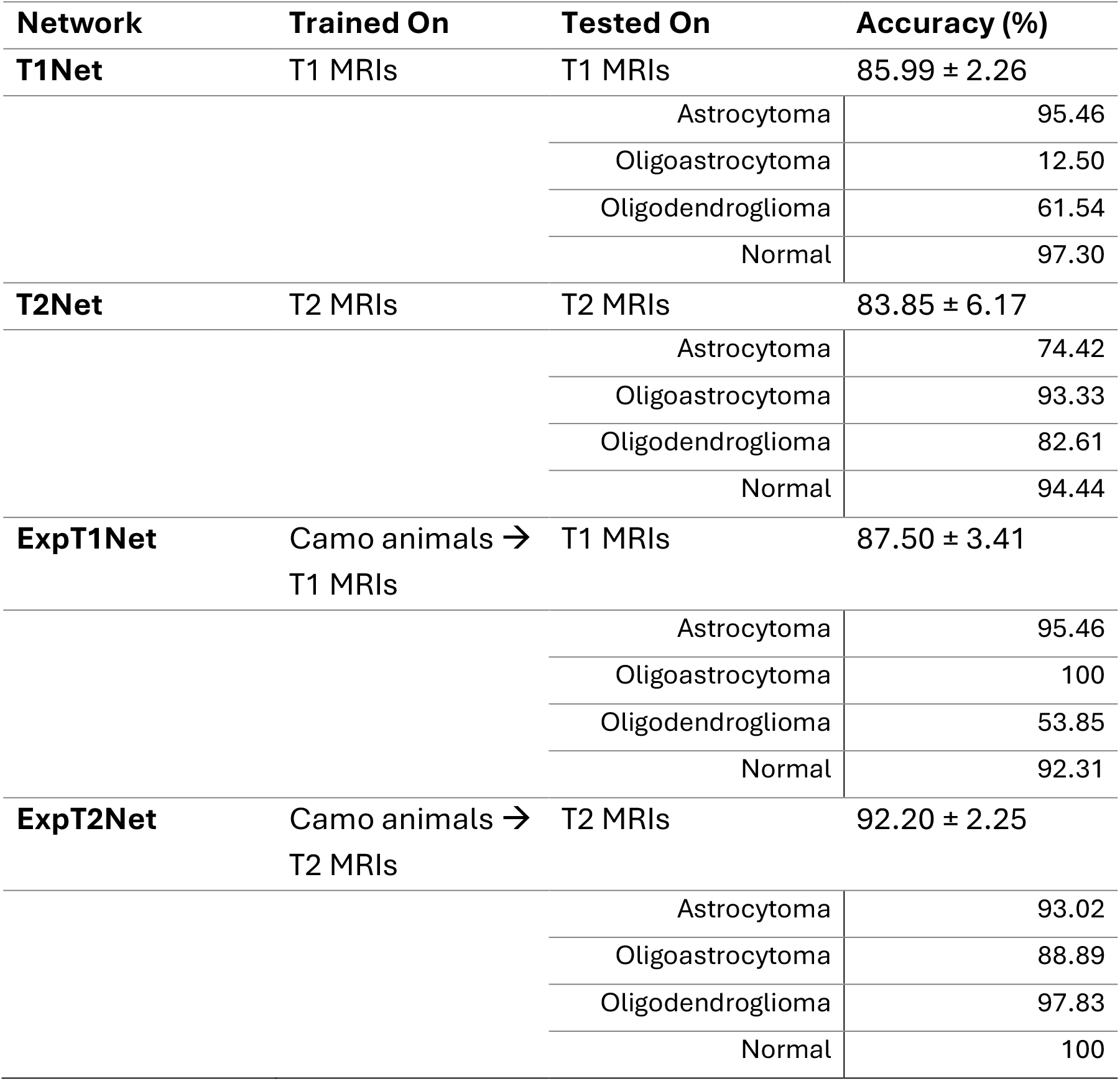
Mean (± SD) model performance across T1- and T2- weighted images, for both seed and transfer trained networks.

The feature spaces of both T1Net and T2Net reveal a distribution of the data in each category relative to another by the networks in a two-dimensional space (Fig 1). We observe a similar pattern of data distribution across both T1Net and T2Net, in which there is a distinct separation between the normal category region and the three glioma category regions on the first principal component. This is consistent with the near-perfect normal image accuracy in both networks mentioned above as it translates to a clear decision boundary between these categories. The three glioma category regions, on the other hand, are blended in both the T1Net and T2Net feature spaces across both the first and second principal components. This represents a more muddled decision boundary between the categories by the networks, again consistent with the lower accuracies for these categories. Despite the glioma categories being mixed together, it is apparent upon closer inspection that the oligoastrocytoma category region mostly occupies the space between the astrocytoma and oligodendroglioma regions in both networks.

### Camouflage Animal Transfer Learning for Tumor Detection

The previously trained camouflage animal detection CNN, ExpCamoNet [10], was used as the seed network for training two transfer trained networks, ExpT1Net and ExpT2Net. Both transfer trained networks achieved a higher average accuracy than their non-transfer trained counterparts (Table 1). The accuracy increase was only significant in the case of ExpT2Net (83.85% to 92.20%, p = .0035), whereas ExpT1Net experienced a much smaller increase in average accuracy (85.99% to 87.5%, p = .3153). The accuracy of the normal category remained near-perfect in the case of ExpT1Net, while ExpT2Net rectified the prior false negatives to achieve perfect accuracy on normal images (compare Figs 1b and 2b). ExpT1Net’s largest improvement was in the oligoastrocytoma category, jumping from 12.50% to 100% (also compare Figs 1a and 2a and the expansion of feature space in ExpT1 Net), with all other categories remaining relatively unchanged. ExpT2Net improved in classification of each category, most notably in its formerly worst astrocytoma category (74.42% to 93.02%, also compare Figs 1b and 2b).

When comparing the feature spaces of ExpT1Net and ExpT2Net after transfer learning to their original versions, we notice changes in data distribution (Fig 2). As expected, the normal category region still occupies an independent region separate from the glioma categories as before, albeit with some noticeable increase in separation on the first principal component between normal and glioma regions indicative of the expansion of feature space in particular for ExpT1 Net. The more obvious transformation is within the glioma categories themselves in the ExpT2Net feature space (Fig 2b), which had the more pronounced quantitative improvement in accuracy. This translates to increased separation of the astrocytoma category region to the point of now showing a much more defined decision boundary, especially when compared to the T2Net feature space (Fig 1b) in which astrocytoma was the worst performing category. This is also true to a lesser extent for the oligoastrocytoma and oligodendroglioma category regions which remained mixed together, but now with more distinct separation between them. The oligoastrocytoma category remains in between the astrocytoma and oligodendroglioma regions as before. These observations hold true for the ExpT1Net feature space, but to a lesser extent since transfer learning was not as beneficial to this network despite the expansion of feature space in ExpT1 Net compared to T1 Net.

### DeepDreamImage Analysis of Trained Networks

DeepDreamImage (DDI) images of each data type for the four trained networks were generated to provide an additional qualitative assessment of the networks’ internal representation of glioma and normal brain MRIs. Figure 3 shows a side-by-side comparison of each DDI for all networks and categories. The seed networks, T1Net and T2Net, appear to produce certain shapes corresponding to the tumor categories they were trained on. For example, the astrocytoma DDI shows a repeating nodal shape pattern throughout the image, whereas the oligodendroglioma DDI shape is more bulbous. The oligoastrocytoma DDI shape falls into an intermediate of the two other glioma categories, appearing to have both a central node and an external bulb shape.

**Figure 3.**
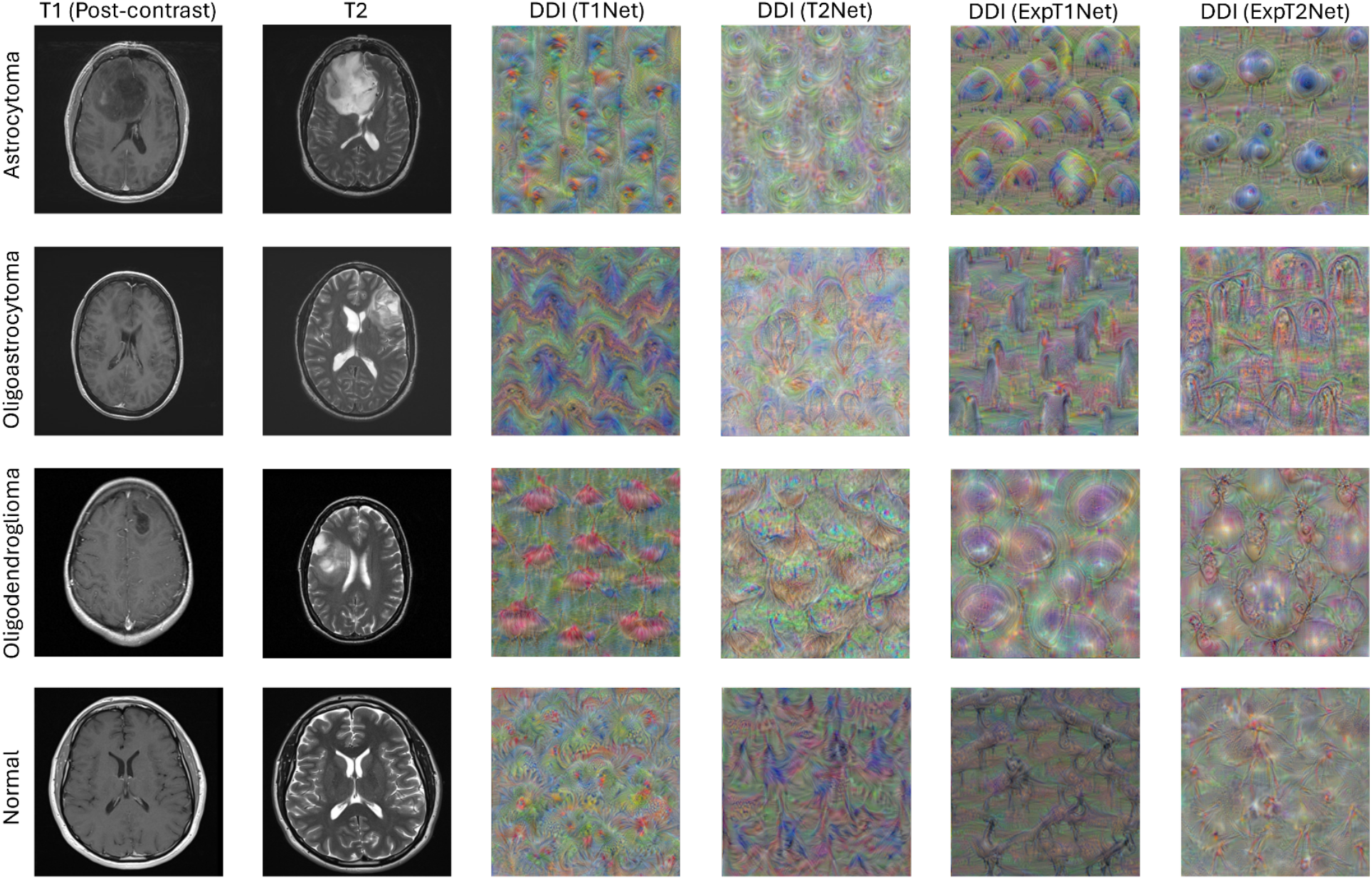
DeepDreamlmage images for T1Net (third column), T2Net (fourth column), ExpTINet (fifth column), and ExpT2Net (sixth column). Example MRIs of both post-contrast T1-weighted (first column) and T2-weighted (second column) are also shown for comparison.

After transfer learning, the ExpT1Net and ExpT2Net DDI generated images have similar patterns that are more pronounced. Clearer and more defined versions of the shapes described in T1Net and T2Net DDI emerge, while retaining their original node/bulb characteristics. This again represents how transfer learning enables the networks to produce more distinct internal representations of the data. For normal image DDIs, on the other hand, they seem to lack any clear shapes like those observed in glioma DDIs. This suggests that the networks associate a lack of distinct features with the normal category classification label, while associating pronounced nodal and bulbous shapes with the various glioma categories.

### ImageSensitivity Analysis

ImageSensitivity maps provide another layer of analysis for identifying salient features used by a network to make a classification decision. Figures 4&5 show the side-by-side comparison of the sensitivity maps across all image categories and networks. The sensitivity maps for T1Net and T2Net show that these networks typically identify the tumor itself as the most salient feature of the image, as one would expect. In the case of T1Net especially, the areas of highest sensitivity are broad and cover a large portion of the brain’s surface (Fig 4). In certain instances (T1 oligoastrocytoma, T1 oligodendroglioma, T2 oligoastrocytoma, and T2 oligodendroglioma) the networks appear to also focus on the tissue adjacent to the tumor but not directly on the tumor itself.

**Figure 4.**
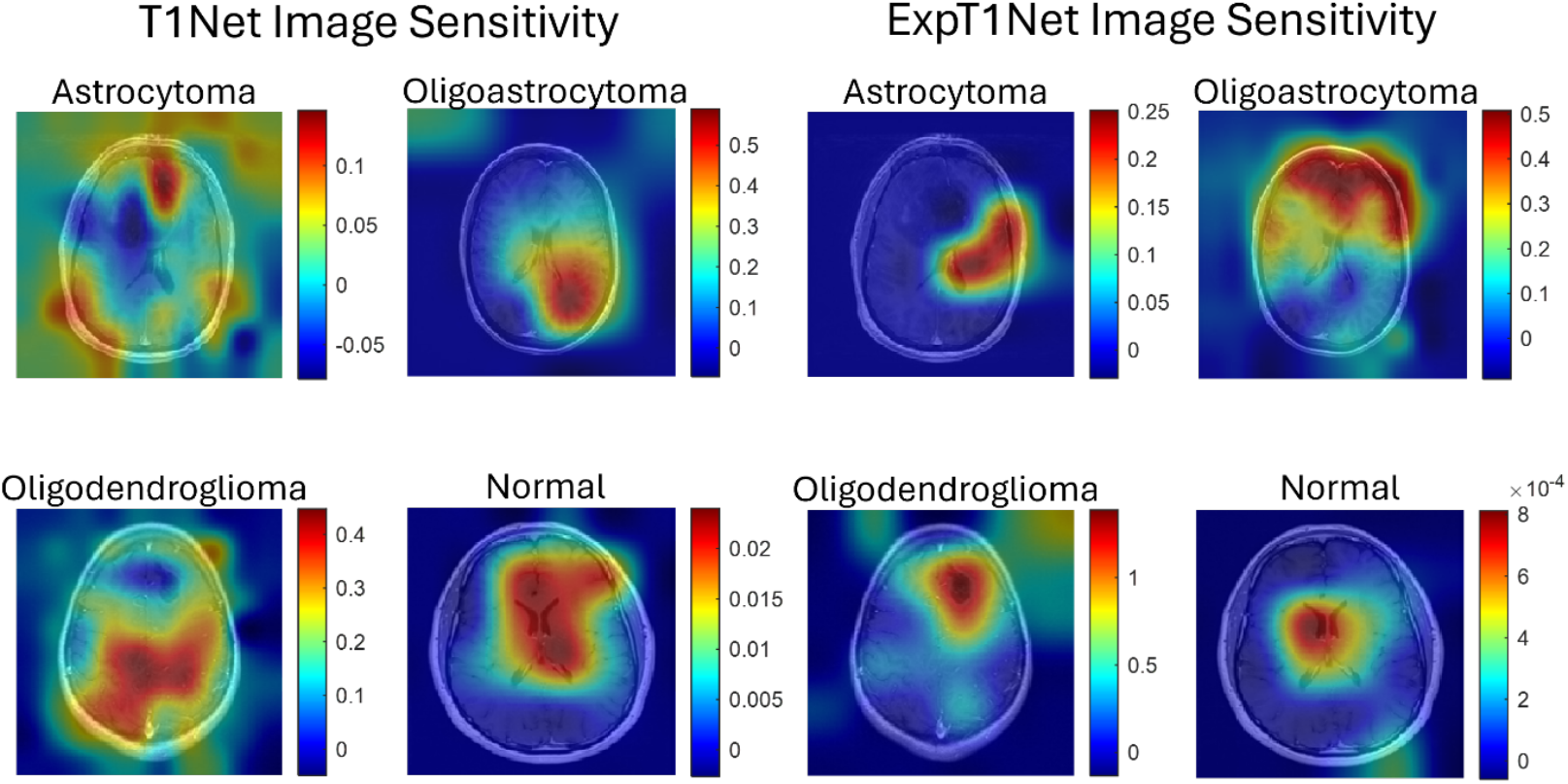
Image sensitivity maps for T1Net and ExpTINet showing transfer learning effect. Sensitivity maps were generated using oS and gC functions, on both softmax and fully connected layers. The underlying sample images are the same as those from Figure 3.

**Figure 5.**
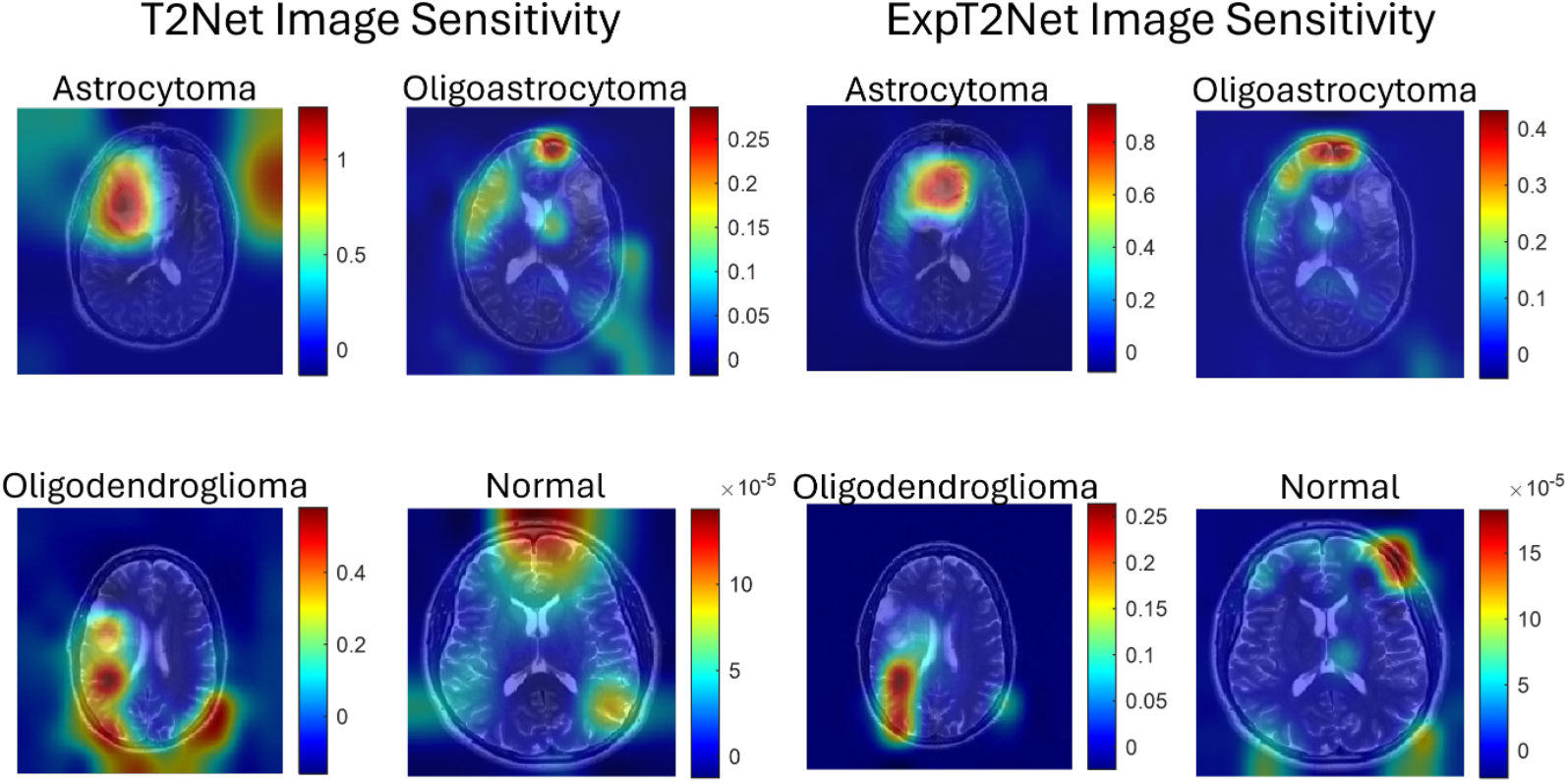
Image sensitivity maps for T2Net and ExpT2Net showing transfer Learning effect. Sensitivity maps were generated using oS and gC functions, on both softmax and fully connected layers. The underlying sample images are the same as those from Figure 3.

After transfer learning, both ExpT1Net and ExpT2Net’s sensitivity maps show a more precise and specific focus on certain features compared to the broader sensitivity in the seed networks. Again, this includes sensitivity to either the tumor itself and/or the tissue around it as it becomes distorted by the presence of the tumor. When looking at the image sensitivity maps of the normal images, for both seed and transfer trained networks, the most salient areas are either in the middle of the brain (in the case of T1Net and ExpT1Net, Fig 4) or bordering with the skull (T2Net and ExpT2Net, Fig 5). The color bar corresponding to each normal image, however, shows that the degree of the network’s sensitivity to these regions is orders of magnitude smaller than in the case of the glioma sensitivity maps. In other words, the regions that appear most sensitive in the normal brain images are in fact not very salient, further reinforcing the previously introduced idea that the networks associate a lack of distinct features with the normal category classification label.

## Discussion

This investigation is the first of its kind to apply animal camouflage transfer learning to deep neural network training on a tumor detection and classification task. Our results demonstrate that this approach to deep neural network training is promising, specifically when using T2-weighted MRI data which showed the greatest improvement in testing accuracy. The effectiveness of this method was demonstrated both quantitatively, with traditional accuracy metrics, and qualitatively through feature spaces, image sensitivity maps, and the DeepDreamImage algorithm. Our methods provide insight into how a camouflage animal training paradigm can transform the internal representation of a neural network on brain cancer imaging data, and visualize how the network’s decision-making process resembles that of a human radiologist.

The feature spaces of the trained networks organized for the glioma data in such a way that the oligoastrocytoma category region occupied the space in between the astrocytoma and oligodendroglioma regions. Even though oligoastrocytomas are no longer recognized as a glioma as they are considered mixed cell tumors, we used the available oligoastrocytoma data in this study to evaluate the networks’ performance with its inclusion. We found that the networks intuitively recognized the mixed cell appearance of oligoastrocytomas as an intermediate between astrocytomas and oligoastrocytomas, and thus organized them as such relative to the two other glioma categories. In other words, the networks’ internal representation of oligoastrocytomas is somehow consistent with the updated classification of gliomas.

The feature spaces also illustrate an increased generalization ability of the neural networks after transfer learning accompanied with expanded region of each category in feature spaces. As described in the Results section, transfer learning led to an increase in mean accuracy for the networks as well as feature space transformations consistent with this increase. The increased separation between category regions represents a more defined decision boundary by the network, which is consistent with an enhanced generalization ability gained by the networks from camouflage animal transfer learning. Since the process of camouflage animal detection itself is heavily reliant on the ability to generalize the appearance of an animal under different conditions, it is possible that this ability was abstracted by the tumor detection networks during the transfer learning process. This would mean that ExpT1Net and ExpT2Net perform higher on their task because they gained the ability to generalize the appearance of the tumors under differing conditions and thus detect features more consistently than before. This is further reinforced by the DeepDreamImage images generated by the transfer trained networks. In these images, we see that the shape the network associates with each tumor category becomes more defined and pronounced, again indicating an enhanced generalization ability to detect these key tumor features across various brain MRIs and classify them accordingly.

The image sensitivity maps show a high-level decision-making process by the networks to detect and classify tumors. In several cases, the networks did not highlight the tumor as the most salient feature. Rather, it focused on the surrounding tissue directly adjacent to the tumor, and even in more distal regions in some cases. This approach of examining the distortion of neighboring tissue caused by the tumor is similar to that of radiologists when studying similar cancerous MRIs [18], which the network determined without prior suggestion. Specifically, the perceived saliency to brain regions directly opposite the tumor itself in some cases suggests an ability of the network to compare symmetrical regions for structural displacements and shifts secondary to tumors, representing a highly sensitive feature of the network. Through training on the glioma data, the network learned to look at the areas affected by the tumor and not just at the tumor itself when making a classification decision.

This study had some limitations. First, the imbalance of the datasets may have affected the networks’ performance on certain categories. The datasets used contained more T2-weighted MRIs than post contrast T1-weighted, which may explain the discrepancy between ExpT1Net and ExpT2Net after transfer learning. Furthermore, the number of images within each category was imbalanced, especially with T1-weighted images. For example, there were more post-contrast T1 astrocytoma MRIs than any other category, which is likely why T1Net performed best on that category relative to the other glioma types. Second, the lack of standardization between images from different dataset sources may have factored into the networks’ performance. The MRIs from TCIA were slightly different in formatting than those obtained from the VA healthcare system, and these differences may have been distracted or used by the network when making a classification decision.

In conclusion, convolutional neural network models were trained on post-contrast T1-weighted and T2-weighted brain MRIs to detect and classify gliomas. Transfer learning from previously trained camouflage animal detection models yielded the highest accuracy on this task, specifically with T2-weighted images where the change in accuracy after transfer learning was significant. The highest performing model achieved perfect accuracy on normal brain MRIs, with no false negatives or positives, and near perfect accuracy on astrocytoma and oligodendroglioma categories. Qualitative analysis of the models’ performance revealed an improved generalization ability following animal transfer learning, which included a distinct, tumor-specific feature-based approach to image classification. These models have been made publicly available for further investigation and potential clinical application in glioma detection and classification from MRI data.

